# Rhesus monkeys use both eye and head gaze to reallocate covert spatial attention facilitating visual perception

**DOI:** 10.1101/2025.07.07.663380

**Authors:** Masih Shafiei, Matthias Reik, Marius Görner, Nick Taubert, Martin Giese, Peter Thier

## Abstract

Non-verbal cues, particularly eye gaze, significantly shape human social interactions. Although non-human primates reliably follow head gaze, their capacity to use eye gaze alone for inferring the other’s focus of attention remains debated. We investigated this question using a realistic rhesus monkey head avatar that directed its gaze toward one of two LEDs (left or right), employing either eye movements alone or combined eye and head movements. After a randomly chosen interval (ranging 50-400 ms) from gaze presentation, one LED transiently increased its luminance to near-threshold levels. Rhesus monkeys were trained to detect and report this luminance change via a saccade to the corresponding LED, independent of the avatar’s gaze direction, to receive rewards. Our results showed that head-gaze cues robustly directed covert attention toward gaze-congruent targets with short delays, indicative of reflex-like, stimulus-driven orienting. In contrast, eye gaze alone, at comparable amplitudes, did not affect attention shifts. However, increasing the avatar’s size and eye-gaze amplitude, simulating a close-range interaction, made eye gaze cues effective in guiding attention. These findings demonstrate that rhesus monkeys possess the capacity to use eye-gaze cues to determine conspecifics’ attentional targets, and validate and underscore the utility of 3D animal models as powerful tools for generating realistic yet precisely controlled stimuli. Our study supports the idea that eye-gaze following is not uniquely human but is an evolutionarily ancient ability, likely shared across Old World monkeys and apes which diverged more than 30 million years ago.

## Introduction

While language serves as the primary channel of communication among humans, nonverbal cues offer a parallel and richly informative modality that leverages additional sensory and motor systems for both signal transmission and reception^1^. In verbal communication, the sender primarily engages articulatory mechanisms, while the receiver relies on the auditory system. In contrast, nonverbal communication recruits the sender’s bodily movements, such as facial expressions, gestures, and posture, and mainly the receiver’s visual system to convey and interpret meaning. This multi-modal interaction enables the receiver to integrate verbal and nonverbal information, thereby enhancing the efficiency, reliability, and perhaps depth of communication^2^. Nonverbal cues also provide a means to assess the congruence and trustworthiness of verbal content, allowing the receiver to cross-reference messages across modalities^3^. Importantly, nonverbal cues are not merely supplementary to language but can operate independently to convey information. For example, an individual can silently draw another’s attention to an object of interest by first establishing eye contact and then shifting their gaze toward the target. This sequence prompts the observer to follow the initiator’s gaze and locate the object, illustrating how attention can be directed to targets through non-verbal cues alone^4^. Among various nonverbal cues, gaze direction, defined by the spatial orientation of the otheŕs eyes, face, or body, serves as a salient indicator of their current focus of attention. The ability to align one’s own attention with the otheŕs gaze, known as gaze following, allows individuals to access relevant information in the environment, such as potential threats or resources that might otherwise go unnoticed^5^. Moreover, gaze following supports higher-order social cognitive processes by enabling individuals to infer others’ mental states, such as intentions, desires, or beliefs, a foundational capacity helping humans to form a Theory of Mind (ToM)^6,7^. In humans, gaze following has been strongly associated with first language acquisition^8–10^, and impairments in this ability have been linked to delays in social development, particularly among individuals with autism spectrum disorder (ASD)^11–14^.

Previous research has explored the evolutionary origins of gaze-following by studying a range of animal species, whose evolutionary trajectories diverged from the human at different times. Non-human primates (NHPs) have been of particular interest given their close evolutionary relationship with humans and their ability to engage in complex social behaviors^15,16^. The goal has been to examine the extent of similarity between the gaze-following abilities of humans and NHPs. Although it has been consistently shown that both humans and NHP follow the gaze direction of either conspecifics or humans to co-orient their attentional focus with the otheŕs, the type of gaze cues used varies across species^17–44^. Humans are capable of accurately following gaze direction in 3D space by exploiting eye direction relative to the head together with information on head and body orientation^4,45,46^. This ability has long been linked to the distinctive morphology of the human eye characterized by a very bright sclera, and a horizontally elongated shape that exposes more of the sclera, features thought to make the eye markedly more conspicuous than that of NHPs^47,48^. However, Recent quantitative work contests this view; when eye visibility is studied with finer metrics such as iris-to-sclera luminance contrast or the proportion of sclera revealed during averted (rather than direct) gaze, human eyes are no longer significantly different from several NHP species^49–52^. On the other hand, although previous reports consistently support NHPs ability to use the other’s head direction to identify their object of interest^17–23,25–28,30–42^, there is still no clear consensus on whether they can follow the others’ eye-gaze in the absence of accompanying head turns^19,21,23,27,33–35,42–44,53^. This lack of agreement may stem from methodological differences across studies. First, several studies have relied on gaze cues from live human demonstrators^19,23,27,33–35,42,44^, which are likely unnatural for NHPs. In others, static images of conspecifics were used^21,53^ instead, but these lack the dynamic motion inherent to natural gaze cues. Second, some studies examined gaze-following responses in non-object-oriented tasks^23,35,42,53^, known to attenuate orienting responses with repeated exposure^32,54^, or linked gaze direction to object detection in a reward-based task^19,20,27,33^, potentially training animals to associate gaze with reward rather than revealing spontaneous or intrinsic gaze-following abilities. Finally, a key question is whether eye direction relative to the head is used by NHPs as a vectorial cue that indicates spatial direction, or merely as a modulatory signal that enhances or interferes with the processing of head-derived information. This distinction is essential when considering whether NHPs’ limited use of eye-gaze cues reflects a perceptual constraint in processing eye gaze or simply poorer input quality, especially when the visibility of the eyes may be reduced. If the latter explanation holds, eye gaze may play a more significant role where its informational value is highest like when social interactions occur at a closer range.

To address these limitations of the previous research, we designed the current study. To overcome the challenge of providing ecologically valid yet controlled gaze cues, we decided to deploy a hyper-realistic rhesus monkey head avatar as demonstrator. This avatar, known to elicit completely natural behavioral reaction in monkey observers^55^, allowed for precise control over gaze direction, as well as control over the relative contributions of head and eye orientation, all while maintaining natural motion dynamics. The use of the avatar enabled us to investigate our core question: whether eyes alone can modulate the observeŕs focus of spatial attention. To this end, we used variants of gaze shifts presented by the avatar: a first one in which the avatar turned his head towards a target with the eyes moving with the head and a second one in which only the eyes were directed to the target whereas the head was kept straight. To address the second limitation of previous work, we designed a paradigm that ensured that gaze-following itself was not reinforced, isolating the possible influence of spontaneous gaze-following behavior on attention and perception. Specifically, we implemented a near-threshold luminance detection task, preceded by a gaze cue. The avatar randomly directed the gaze to one of the two LEDs, placed to the right and left. After a variable time from the gaze cue, one of the LEDs randomly increased its luminance to a near-threshold level for a brief duration. The rhesus monkeys were trained to detect the change and report it, regardless of the avatar’s gaze direction, by making a saccade to the corresponding LED to receive reward for correct responses. The assumption was that gaze induced shifts of covert attention should manifest themselves in an improvement of luminance change detection thresholds. Moreover, by determining the shortest stimulus onset asynchrony (SOA), i.e. the time between the onset of the gaze cue and the time of the luminance change, we were allowed to assess whether a shift of covert attention occurs fast enough to qualify as an exogenous, stimulus-driven response. We show that head gaze cues draw the observeŕs focus of attention to objects of interest with short delays which are fully compatible with reflex-like stimulus-driven behavior, but not so for eye gaze shifts of the same amplitude in the absence of accompanying head movements. However, also eye gaze shifts become effective if the distance to the avatar head is substantially reduced and the amplitude of the eye gaze increased, simulating close-range social interactions with improved eye visibility.

## Methods

All experimental procedures complied with German and European Guidelines for the Care and Use of Laboratory Animals and were reviewed and approved by the responsible local authorities (Regierungspräsidium Tübingen and Landratsamt Tübingen).

### Experimental subjects

The selection of rhesus monkeys (*Macaca mulatta*) as the animal model for this study was guided by the aim to explore the evolutionary origins of human gaze-following by exploring the gaze-following capabilities of rhesus monkeys, arguably the best-explored representative of old world monkeys whose visual and visuo-motor systems match those of humans to a large extent^56^. We used two male monkeys, referred to as “monkey E” and “monkey C”. Monkey E was naïve to tasks resembling those used in the present study. Monkey C had previously undergone partial training for an unrelated project involving static images of conspecific head-gaze cues, where he was required to follow or ignore the gaze cue based on a rule dictated by a pre-cue. However, his training was not completed due to unrelated circumstances. Both animals had existing headpost implants, which enabled the painless fixation of the head necessary for high-precision eye-tracking (for a description of the detailed neurosurgical procedures, see Prsa, et. al. 2011^57^). Additionally, monkey E had received a scleral search coil implant, enabling eye tracking using an in-house system developed based on a design originally described by Bechert and Koenig (1996)^58^. Eye movements of monkey C were tracked via high-resolution video tracking using the EyeLink 1000 Plus desktop system (SR Research Ltd., Canada).

On experimental days, animals were maintained under controlled water access protocols, that ensured that water was received as a positive reinforcement upon successful trial completion. To enhance motivation, monkey C was occasionally rewarded with apple juice instead of water because, unlike monkey E, he was very fond of juice.

### Experimental setup

Figure 1A illustrates a schematic of the experimental setup. Head-fixed animals, calmly seated in a primate chair, were positioned inside a dimly lit, sound-attenuated booth. A 24-inch LCD monitor (BenQ XL2411-B; 1920 × 1080 resolution; 60 Hz refresh rate) was mounted 60 cm in front of the animals, centered on their neutral line of sight. A transparent plexiglass panel was placed 14 cm in front of the screen. Two LEDs were fixed on this panel at 7.5 cm eccentricity to the left and right of the center along the horizontal axis. The LEDs were illuminated at a baseline luminance of 0.09 cd/m² for monkey E and 0.01 cd/m² for monkey C.

**Figure 1.**
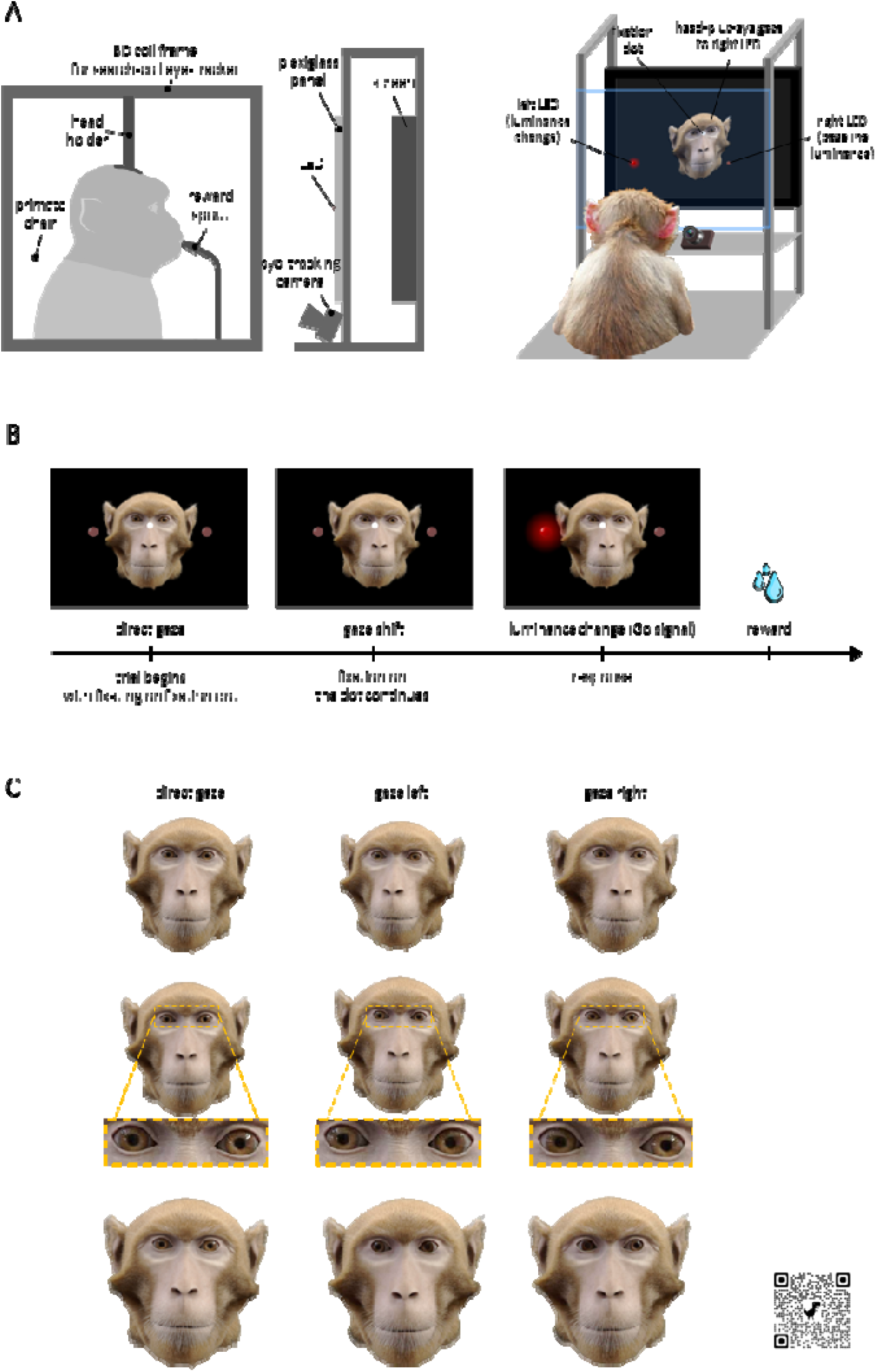
(A) Schematic of the experimental setup. The coil frame used for search-coil eye tracking, shown on the left panel, was employed exclusively for monkey E. On the right, a simplified illustration of the experimental setup is shown; components such as the head-fixation apparatus, reward delivery system, and primate chair have been omitted for clarity. (B) Sequence of events in a trial. This panel shows an incongruent trial, where the avatar’s gaze is directed towards the distractor LED. (C) Each row represents a different gaze-cue type: head gaze (top), small-amplitude eye gaze (middle), and large-amplitude eye gaze (bottom). Columns show video snapshots depicting, from left to right: a direct gaze, a gaze toward the left LED, and a gaze toward the right LED. Insets in the middle row provide a zoomed-in view of the peri-orbital region to enhance visualization of the gaze direction. Videos of the different gaze cues are accessible via the QR code in panel C, or through the following link: https://doi.org/10.6084/m9.figshare.28769981.

Rewards were delivered directly to the animal’s mouth via a spout positioned in front of him, controlled by a fully automated, in-house developed computer-controlled reward system.

### Behavioral paradigm

Figure 1 illustrates the trial structure and the different gaze cue conditions. At the start of each experimental session, eye movements were calibrated using a standard nine-target positions calibration paradigm. The LEDs were kept off during the calibration. Each trial in the main behavioral paradigm, designed to elicit gaze-following, began with the appearance of a forward-facing avatar, with a white fixation dot (0.2° diameter; CF=central fixation dot) on its nose bridge as shown in Figure 1A and B. This dot was horizontally aligned midway between the two LEDs. To initiate a trial, monkeys were required to maintain fixation within an invisible 5° × 5° window around the CF for 500 ms. Following successful fixation, the avatar shifted its gaze randomly toward one of the two LEDs. After a variable delay, referred to as the stimulus onset asynchrony (SOA), one of the LEDs’ luminance increased briefly (for 300 ms), identifying the target LED. The target LED was chosen at random, independent of the avatar’s gaze direction. Its luminance rose to 0.28 cd/m² (from 0.09 cd/m²) for monkey E and 0.07 cd/m² (from 0.01 cd/m²) for monkey C yielding ΔLum (delta luminance) values of 0.18 and 0.06, respectively. These individualized ΔLum values were determined for each animal based on a perceptual threshold estimation procedure that identified the luminance change required for 75% detection accuracy (see Methods section “Perceptual threshold estimation”). The onset of the ΔLum stimulus defined the earliest time the monkey could report the target LED by making a saccade towards it, regardless of the avatar’s gaze direction. Correct responses were rewarded, while incorrect choices (saccades made to the distractor LED) were not. After either a correct or incorrect response, an inter-trial interval (ITI) ranging between 700 to 1500 ms was introduced, during which the avatar’s gaze returned to its neutral position. Trials were self-paced allowing monkeys to initiate the next trial by fixating on the CF.

If a monkey broke fixation after the gaze cue but before ΔLum, the same trial was repeated. Trials were aborted if the monkey failed to respond within 1.5 s of the avatar’s gaze shift. To help monkey C differentiate between premature responses and incorrect choices, an aversive buzz tone was played when the monkey responded too early, before the ΔLum appeared. Monkey E, whose data collection was completed earlier and who showed no difficulty in making this distinction, did not require the use of the buzz tone (see the ’Animal training’ section for more details).

To examine the temporal dynamics of attention, we tested five different SOAs: 50, 100, 200, 300, and 400 ms. In each session, a subset of two or three SOAs was selected pseudo-randomly, and one SOA from the subset was randomly assigned to a given trial. Three distinct gaze cue types were tested (directed either left or right): eye gaze accompanied by head turns (figure 1C, first row), eye gaze without head movement (figure 1C, second row), large-amplitude eye gaze without head movement, featuring a larger avatar head (figure 1C, third row). Only one gaze cue type was presented per session. Videos of the gaze stimuli can be accessed via the QR code or URL provided in figure 1. In monkey C, under the eye-only gaze condition prevailing in experiment 1, we considered three SOAs (50, 200, and 400 ms), expected to encompass the SOAs most probably promising significant effects based on the findings on head-gaze following. We intended to test the remaining two SOAs (100 and 300 ms) examined in the head-gaze variant of the experiment in case the results obtained would necessitate further refinement, which turned out unnecessary.

In experiment 1, we compared responses to the avatar making 7.7° gaze shift by either moving eyes and head together with the eyes kept straight relative to the head (“head-gaze”) or only the eyes (“eye gaze”) while the head stayed put (figure 1C). In experiment 2, we realized a more salient eye-gaze cue by enlarging the avatar size by 40%, while maintaining all other experimental parameters constant. This manipulation resulted in an increased amplitude of gaze deviation, reaching 10.8° to either side. While monkey E was able to cope with this configuration, monkey C frequently broke fixation before even the gaze cue was presented, forcing us not to consider this monkey any further in this experiment. We hypothesize that the larger avatar head in this condition may have been experienced as too dominant and threatening, overloading the relatively small monkey C. In support of this interpretation, we observed that monkey C immediately resumed task engagement and fixated successfully upon switching the avatar back to its original size in which the head appeared smaller again. Note that we did not consider collecting responses to larger eye gaze shifts for this latter configuration as it would have been required to move the LEDs further out on the transparent panel.

### Animal training

The monkeys were initially trained to associate an increase in LED luminance from the fixed baseline luminance value with a reward. This luminance change (ΔLum) was randomly presented by one of two LEDs. Task difficulty gradually increased by reducing the magnitude and duration of the ΔLum. Once the monkeys consistently associated brief changes in LED luminance, regardless of absolute luminance levels or LED position, with a reward, they progressed to the next phase of the experiment, “perceptual threshold assessment”.

### Perceptual threshold assessment

This phase of the study aimed to determine the ΔLum corresponding to the individual point of subjective equality (PSE), defined as the luminance change resulting in an average 25% increase in correct responses, relative to the baseline chance level of 50%. To achieve this, we systematically varied ΔLum, while maintaining the duration of luminance change constant at 300 ms, enabling an assessment of the relationship between ΔLum and detection accuracy. A simplified version of the main behavioral paradigm was employed during this phase. To simulate the light emitted by the avatar in the main experiment, a colored silhouette of the avatar, matched in luminance, was displayed throughout the perceptual threshold assessment trials (see supplementary figure 1A).

Each trial began with the appearance of a CF superimposed on the silhouette. The monkey was required to fixate on the CF for the luminance change event to occur. In monkey E, the luminance changes of the LED occurred after a variable delay, randomly chosen from 50, 100, 200 ms, following the CF presentation. In contrast, monkey C experienced a fixed temporal delay of 300 ms between the CF presentation and the luminance change. In both monkeys, the luminance change lasted 300 ms and appeared randomly at one of two possible LEDs. Following the luminance change, the monkeys had a maximum of 1500 ms to initiate a saccadic response. The ΔLum values ranged from high levels, yielding near-perfect detection accuracy, to low levels, resulting in a detection accuracy no better than chance. Correct responses were rewarded with immediate positive reinforcement (a drop of water). An inter-trial interval (ITI) of 700-1500 ms separated trials.

The ΔLum values corresponding to the PSE were 0.18 cd/m² for monkey E and 0.06 cd/m² for monkey C, which were subsequently used in the main experiment (see supplementary figure 1B). The observed differences in PSE between subjects likely reflect individual variations in their sensitivity to perceive changes in luminance, e.g. due non-corrected ametropia, mild enough to not interfere with sufficiently precise fixation and saccades.

### Data processing and statistical analysis

Data preprocessing was conducted using a custom-written script in Julia (version 1.10.3). Statistical data analysis and visualization were performed in R (version 4.4.2, 2024-10-31 ucrt)^59^ using the following packages: *ggplot2* (v3.5.1)^60^, *lme4* (v1.1.36)^61^, *car* (v3.1.3)^62^, *emmeans* (v1.10.7)^63^, and *performance* (v0.13.0)^64^. Estimation of ΔLum values, yielding an average detection accuracy of 75% was carried out in MATLAB (R2023a)^65^ using the *Psignifit* toolbox (v4). Hit rate for each given ΔLum level were modelled using a logistic psychometric function, with the lower asymptote fixed at 50% to reflect chance-level performance.

Eye-tracking data, recorded as a time series of horizontal eye positions, was smoothed using a Savitzky-Golay filter (11 ms window, first-order polynomial). The first derivative of the smoothed eye position data was computed to extract the onset and direction of the response saccade. Reaction time (RT) was defined as the temporal difference between ΔLum onset and saccade onset. Trials with reaction times below 70 ms or above 1500 ms were excluded, as they were considered unrelated to the task. This resulted in the removal of 5.7% of trials for monkey E (5.3% in the head-gaze condition, 5.9% in the small-amplitude eye-gaze, and 2.6% in the large-amplitude eye-gaze) and 10.7% for monkey C (11.7% in the head-gaze, and 9.7% in the small-amplitude eye-gaze). The final dataset comprised of 9,717 head-gaze trials, 35,830 small-amplitude eye-gaze trials, and 26,885 large-amplitude eye-gaze trials for monkey E. For monkey C, 10,154 head-gaze trials and 10,820 small-amplitude eye-gaze were retained for analysis.

To assess the impact of gaze cue congruency on detection accuracy, generalized linear mixed models (GLMMs) with a logistic link function and maximum likelihood estimation were used. The dependent variable was *hit*, coded as a binary outcome (1 = correct, 0 = incorrect). Fixed predictors were *congruency* (congruent vs. incongruent), *SOA* (50, 100, 200, 300, and 400 ms), *target direction* (left and right), normalized log-transformed *RT*, *SOA × congruency*, *SOA × target direction*, and *SOA* × *normalized log-transformed RT*. Reaction times were log-transformed and z-scored prior to analysis. For Monkey C, only three SOAs (50, 200, and 400 ms) were used in the eye-only condition as noted earlier. Target direction was specifically included to detect potential lateralized biases in performance. The GLMM estimates coefficients for fixed predictors in log-odds units, representing the log odds ratio (OR) of a correct detection given a one-unit change in a continuous predictor or a comparison between a given and reference level for categorical predictors. Where appropriate, log-OR values were converted to odds ratios (ORs) using the exponential function to facilitate interpretation.

Model selection followed a backward stepwise elimination approach using data from the first subject (monkey E) as an exploratory dataset. At each iteration, predictors were removed if their p-values were ≥ 0.05 and/or, for predictors with 1 degree of freedom (df), if their Generalized Variance Inflation Factor (GVIF) exceeded 3.2^62,66^. After each elimination, the model was re-estimated and re-evaluated using the same criteria. This process generated a series of candidate models, which were compared using the Akaike Information Criterion (AIC) ^67^. The model with the lowest AIC, provided all retained predictors with df = 1 had GVIF < 3.2, was selected as the best-fitting model. A ΔAIC > 10 was considered strong evidence in favor of the model with the lower AIC^68^. The significance of the random effect was evaluated by comparing the final GLMM to an equivalent linear model (LM) without the random term, based on AIC. In addition, the 95% confidence interval (CI) for the variance explained by the random effect was computed; the random effect was retained only if the CI did not include zero^69,70^. We also report the adjusted intraclass correlation coefficient (ICC) for the random factor in the best-fitting GLMMs to quantify the proportion of residual variance attributed to session-level effects after accounting for fixed predictors. The final models derived from Monkey E were then applied to data from Monkey C to assess generalizability. Consistency in the direction and statistical significance of key predictors across both subjects was interpreted as evidence of model robustness and replicability.

To examine the influence of congruency on hit rates within each SOA level, post hoc pairwise comparisons between congruent and incongruent conditions were performed using the estimated marginal means (EMMs) derived from the best-fitting models. P-values from these comparisons were corrected for multiple testing using the *Benjamini-Hochberg false discovery rate* (FDR) procedure^71^. Additionally, to evaluate the overall contribution of each fixed effect in the final models, *Type III Wald chi-square tests* were conducted using analysis of deviance (Type III ANOVA)^72,73^. *P-values* from these tests were also corrected for multiple comparisons using the Benjamini-Hochberg method. We report the *Wald* χ*² statistic* for each fixed effect, which quantifies the extent to which the inclusion of a specific predictor improves model fit, conditional on all other variables in the model.

## Results

### Effect of head-gaze on detection accuracy

We found a significant main effect for congruency, SOA, and target location, along with significant interactions between SOA and target location, as well as SOA and RT, on hit probability in both monkeys (table 1 and figure 2). *Post hoc* pairwise comparisons revealed a significantly higher hit probability in the congruent condition compared to the incongruent condition at SOAs of 50, 100, and 200 ms in both monkeys (see table 2 and figures 2 and 3). This finding suggests that target detection is significantly enhanced when it is congruent with the preceding head gaze cue, provided the cue-target interval does not exceed 200 ms. The odds ratios (ORs) indicated that the likelihood of a hit in the congruent condition was 1.4 to 1.7 times higher in monkey E and 1.25 to 1.4 times higher in monkey C, depending on the SOA (see table 2 and figure 2). Furthermore, at an SOA of 300 ms, monkey C exhibited a significant inverse congruency effect, with hit probability being 1.14 times higher in the incongruent condition than in the congruent condition, suggesting a potential *inhibition of return* effect.

**Figure 2.**
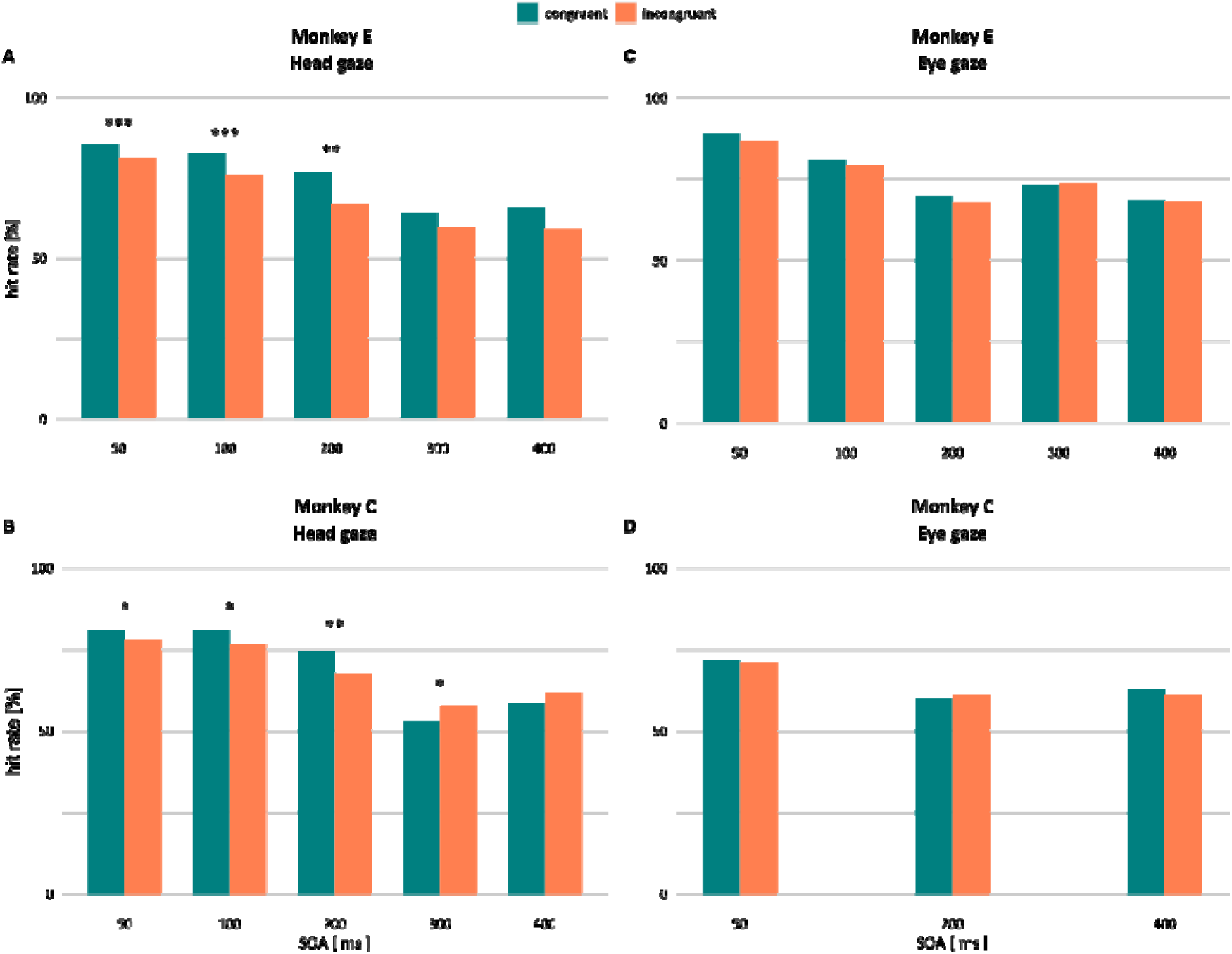
Hit rate as a function of SOA and congruency. Panels (A) and (B) show results from the eye-head condition, while panels (C) and (D) display results from the small-amplitude eye-only condition. Data for Monkey E are presented in the top row (A, C), and data for Monkey C in the bottom row (B, D). A significant effect of congruency on the hit rate is observed in the eye-head condition but not in the eye-only condition. This pattern is consistent across both monkeys. *: p<0.05; **: p<0.01; ***: p<0.001.

**Figure 3.**
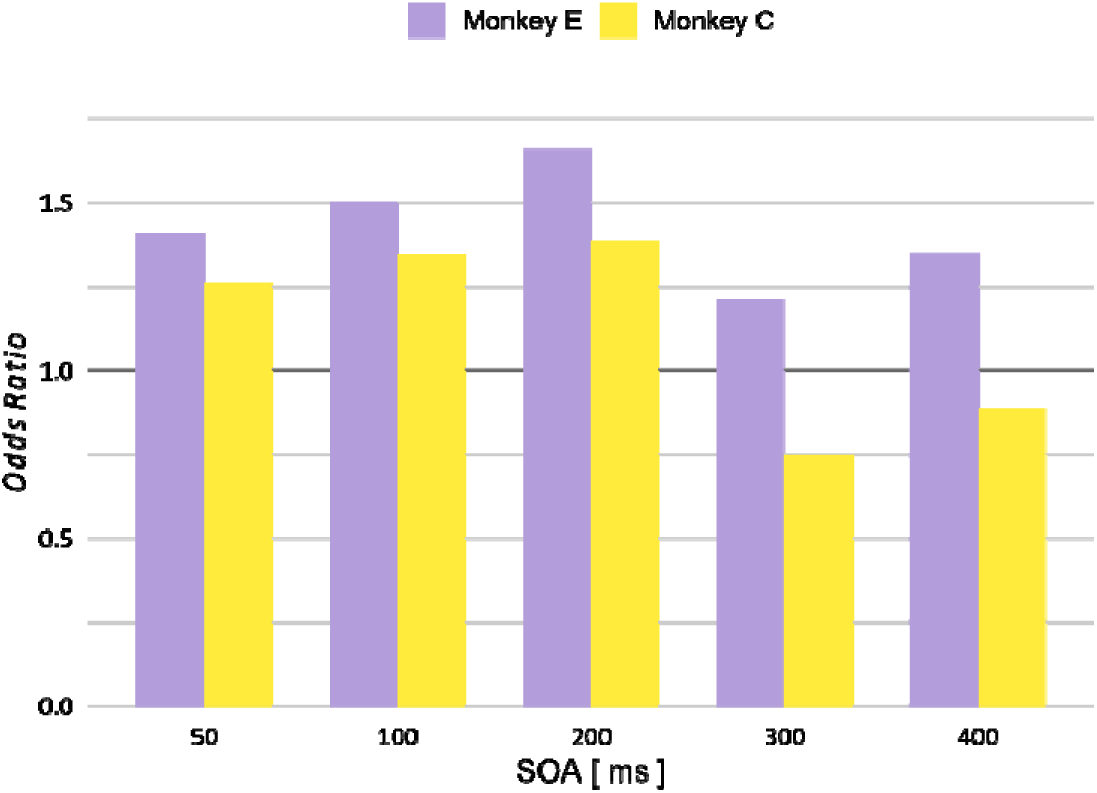
Odds ratios for the effect of congruency in the head-gaze condition on hit probability as a function of SOA, shown separately for each monkey. In both monkeys, the probability of a hit in the congruent condition increases with SOA up to 200 ms. However, a notable decline in the odds ratio is observed for larger SOAs. This decline is more pronounced in monkey C, where the odds ratio drops below 1.0 at 300 ms, consistent with an *inhibition of return* effect.

**Table 1.**
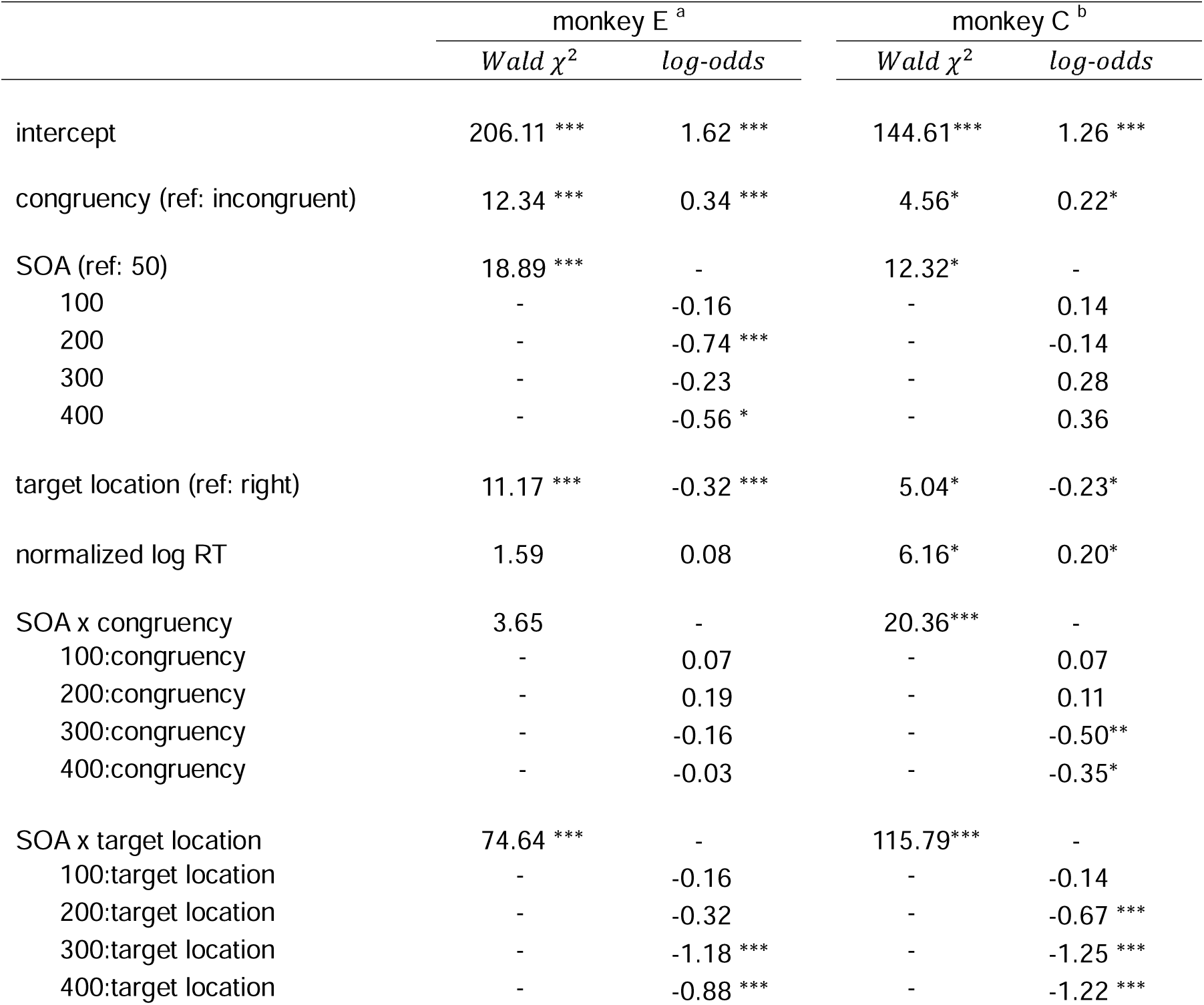
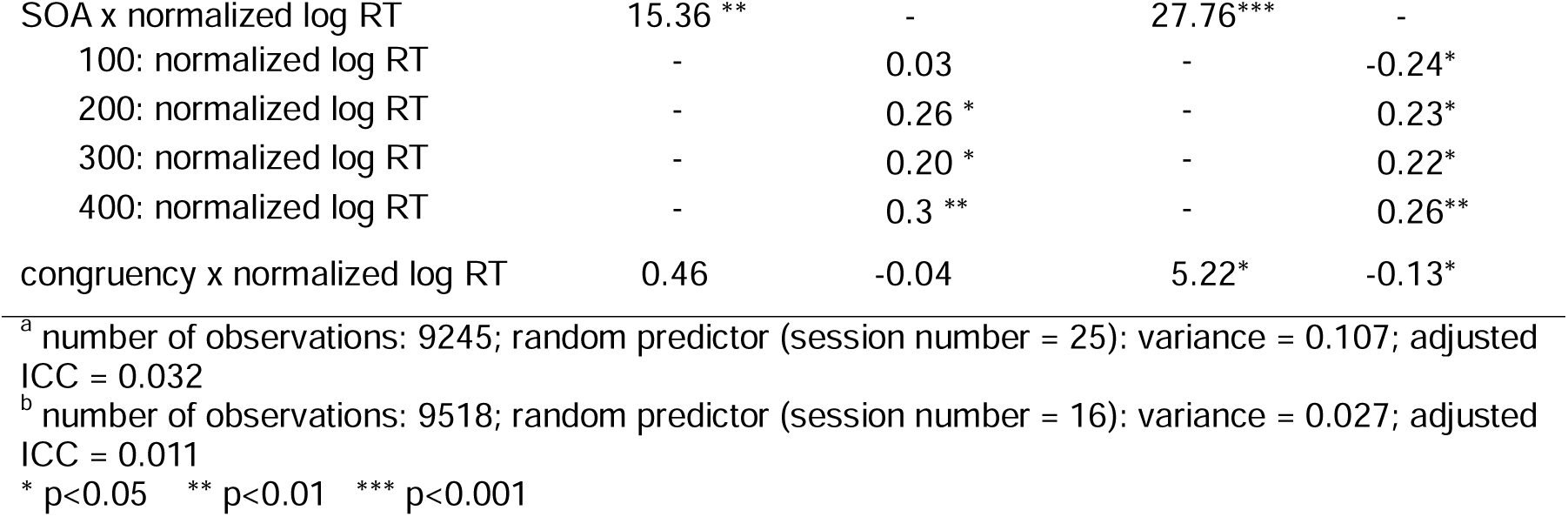
presents the GLMM and ANOVA outputs for the fixed predictors in the eye-head gaze condition.

|  | monkey E <sup>a</sup> |  | monkey C <sup>b</sup> |  |
| --- | --- | --- | --- | --- |
|  | *Wald $\chi^2$* | *log-odds* | *Wald $\chi^2$* | *log-odds* |
| intercept | 206.11 *** | 1.62 *** | 144.61*** | 1.26 *** |
| congruency (ref: incongruent) | 12.34 *** | 0.34 *** | 4.56* | 0.22* |
| SOA (ref: 50) | 18.89 *** | - | 12.32* | - |
| 100 | - | -0.16 | - | 0.14 |
| 200 | - | -0.74 *** | - | -0.14 |
| 300 | - | -0.23 | - | 0.28 |
| 400 | - | -0.56 * | - | 0.36 |
| target location (ref: right) | 11.17 *** | -0.32 *** | 5.04* | -0.23* |
| normalized log RT | 1.59 | 0.08 | 6.16* | 0.20* |
| SOA x congruency | 3.65 | - | 20.36*** | - |
| 100:congruency | - | 0.07 | - | 0.07 |
| 200:congruency | - | 0.19 | - | 0.11 |
| 300:congruency | - | -0.16 | - | -0.50** |
| 400:congruency | - | -0.03 | - | -0.35* |
| SOA x target location | 74.64 *** | - | 115.79*** | - |
| 100:target location | - | -0.16 | - | -0.14 |
| 200:target location | - | -0.32 | - | -0.67 *** |
| 300:target location | - | -1.18 *** | - | -1.25 *** |
| 400:target location | - | -0.88 *** | - | -1.22 *** |
| SOA x normalized log RT | 15.36 ** | - | 27.76*** | - |
| 100: normalized log RT | - | 0.03 | - | -0.24* |
| 200: normalized log RT | - | 0.26 * | - | 0.23* |
| 300: normalized log RT | - | 0.20 * | - | 0.22* |
| 400: normalized log RT | - | 0.3 ** | - | 0.26** |
| congruency x normalized log RT | 0.46 | -0.04 | 5.22* | -0.13* |

<sup>a</sup> number of observations: 9245; random predictor (session number = 25): variance = 0.107; adjusted ICC = 0.032
<sup>b</sup> number of observations: 9518; random predictor (session number = 16): variance = 0.027; adjusted ICC = 0.011
\* p<0.05 \*\* p<0.01 \*\*\* p<0.001

**Table 2.** Odds ratios (OR) and corresponding p-values for *post hoc* comparisons between the congruent and incongruent conditions.

| SOA | monkey E |  |  |  | monkey C |  |  |  |
| --- | --- | --- | --- | --- | --- | --- | --- | --- |
|  | Eye-head |  | Eye-only |  | Eye-head |  | Eye-only |  |
| | OR | $p_{adj}$ | OR | $p_{adj}$ | OR | $p_{adj}$ | OR | $p_{adj}$ |
| 50 | **1.41** | 0.001 | 1.22 | 0.072 | **1.25** | 0.041 | 1.04 | 0.607 |
| 100 | **1.52** | <0.001 | 1.13 | 0.072 | **1.34** | 0.022 | - | - |
| 200 | **1.70** | <0.001 | 1.09 | 0.124 | **1.39** | 0.006 | 0.91 | 0.493 |
| 300 | 1.20 | 0.151 | 1.04 | 0.659 | **0.75** | 0.022 | - | - |
| 400 | 1.36 | 0.151 | 0.99 | 0.793 | 0.88 | 0.291 | 1.09 | 0.793 |

Note. Odds ratios (OR) greater than 1 indicate a higher probability of a hit in the congruent condition compared to the incongruent condition. Bolded values highlight significant ORs for improved readability. $p_{adj}$ : adjusted p-values.

**Table 3.** displays the GLMM and ANOVA outputs for the fixed predictors in the eye-only gaze condition.

|  | monkey E <sup>a</sup> |  | monkey C <sup>b</sup> |  |
| --- | --- | --- | --- | --- |
|  | *Wald $\chi^2$* | *log-odds* | *Wald $\chi^2$* | *log-odds* |
| intercept | 337.15 *** | 1.95 *** | 170.17 *** | 0.92*** |
| congruency (ref: incongreunt) | 5.13 * | 0.34 *** | 0.27 | 0.04 |
| SOA (ref: 50) | 66.33 *** | - | 0.31 | - |
| 100 | - | -0.50 *** | - | - |
| 200 | - | -0.78 *** | - | 0.01 |
| 300 | - | -0.61 *** | - | - |
| 400 | - | -0.85 *** | - | -0.05 |
| SOA x congruency | 7.07 | - | 2.00 | - |
| 100:congruency | - | -0.08 | - | - |
| 200:congruency | - | -0.12 | - | -0.13 |
| 300:congruency | - | -0.24 * | - | - |
| 400:congruency | - | -0.19 | - | 0.05 |
| SOA x target location | 291.26 *** | - | 179.84 *** | - |
| 100:target location | - | -0.20 *** | - | - |
| 200:target location | - | -0.56 *** | - | -0.64*** |
| 300:target location | - | -0.46 *** | - | - |
| 400:target location | - | -0.47 *** | - | -0.83*** |
| SOA x normalized log RT | 795.14 *** | - | 264.95 *** | - |
| 100: normalized log RT | - | 0.32 *** | - | - |
| 200: normalized log RT | - | 0.40 *** | - | 0.41*** |
| 300: normalized log RT | - | 0.40 *** | - | - |
| 400: normalized log RT | - | 0.35 *** | - | 0.39*** |

<sup>a</sup> number of observations: 33751; random predictor (session number = 63): variance = 0.047; adjusted ICC = 0.014
<sup>b</sup> number of observations: 10202; random predictor (session number = 13): variance = 0.012; adjusted ICC = 0.004
\* p<0.05 \*\* p<0.01 \*\*\* p<0.001

Regarding reaction time, the analysis revealed that faster RTs were associated with lower hit probability: a one SD decrease in normalized, log-transformed RT (≈ 624 ms faster) lowered the odds of a correct response by 23.3% (22.8% in monkey E, and 23.7% in monkey C). This effect was moderated by congruency and SOA. Given the temporal uncertainty introduced by variable SOAs, and the low salience of the luminance transient (reduced intensity and brief presentation), hit probability increased with longer response latencies, likely allowing additional processing before the response was made. However, in congruent trials, the reduction in hit probability associated with faster RTs was significantly attenuated; for Monkey C, the decrease in hit probability was reduced by approximately 14% at shorter SOAs (≤ 200 ms). Monkey E showed a similar pattern, with a non-significant reduction of about 25% across all SOAs. Thus, congruency mitigated, but not eliminated, the negative RT-hit association. In congruent trials, the gaze cue provides supplementary spatial information that supported target detection. The covert shift of attention triggered by the gaze direction increased the probability that the brief luminance change occurs within the attentional spotlight. Consequently, faster RTs in these trials are likely affected by facilitated detection of the luminance change besides the random response variability, altering the relationship between RT and hit probability.

The models further revealed that hit probability was systematically influenced by a target direction bias (table 1). Specifically, the probability of a hit was significantly lower for leftward targets compared to rightward targets. This effect varied across SOAs due to significant interactions between target location and SOA. In both monkeys, the most pronounced direction bias was observed at SOA 50 ms (OR = 0.73 in monkey E; OR = 0.80 in monkey C), while the weakest bias occurred at SOA 300 ms (OR = 0.22 in monkey E; OR = 0.23 in monkey C). The consistent rightward bias across both monkeys, along with the comparable range of effect sizes, suggests a shared underlying mechanism influencing response patterns. To ensure that the observed effect of congruency on hit rate was not confounded by target direction bias, we compared the effect size of congruency between the best-fitting GLMM model and a reduced model that excluded target direction as a predictor. Both models exhibited identical fit statistics (AIC = 9372, BIC = 9528.9). Importantly, the fixed effect estimate for congruency remained virtually unchanged between the full and reduced models (β = 0.3426662 vs. 0.3426667), corresponding to a difference in odds ratio of less than 2 × 10□□. This confirms that target direction bias did not meaningfully influence the relationship between congruency and hit rate.

### Effect of small amplitude eye-gaze on detection accuracy

We found significant main effects of congruency and SOA on hit probability in monkey E under the small-amplitude eye-only condition. In contrast, no significant main effects were observed in Monkey C. In both monkeys, significant interaction effects were detected between SOA and target location, as well as between SOA and normalized log-transformed RT. However, *post hoc* comparisons did not reveal any significant differences in hit probability between the congruent and incongruent conditions across SOAs in either monkey (table 2 and figure 2).

### Effect of large amplitude eye-gaze on detection accuracy

So far, our findings suggest that the type of gaze cue offered by the others (head versus eyes-only) determines whether visual spatial attention of monkeys shifted to the target. Specifically, it appears that monkeys are able to use directional information provided by a conspecific’s head-gaze but not by their eye-gaze. However, is this conclusion fully justified? Rather than reflecting a categorical difference, the inefficacy of eye gaze might be secondary to poorer visibility of the small-amplitude eye-gaze, after all eyes cover a much smaller area of the visual field of the observers’ eyes than the face or head. This difference in size might affect how salient eyes are and therefore entail different potentials to guide attention. To assess whether the absence of an eye-gaze-dependent boost in luminance change detection might be a consequence of the low salience of the eye-gaze cue rather than a perceptual constraint in following eye-gaze, we introduced the large-amplitude eye-gaze cue condition. This involved increasing the avatar’s size and the angular deviation of its gaze. All other aspects of the paradigm and experimental setup held constant to isolate the effect of the modified eye-gaze cue on attention modulation.

Using the large-amplitude eye-gaze cue, we observed a significant main effect of congruency, SOA, target location, and notable interactions between SOA and target location, as well as between SOA and normalized log-transformed RT on the probability of a hit (table 4 and figure 4). *Post hoc* pairwise comparisons revealed significantly higher hit probabilities in the congruent condition at SOAs of 50 ms (OR = 1.17, adj. p = 0.04), 100 ms (OR = 1.18, adj. p = 0.04), and 400 ms (OR = 1.30, adj. p = 0.002). These findings strongly suggest that the absence of a congruency effect on hit probability observed with the original eye-gaze cue can be attributed to the lack of conspicuity of the gaze signal. By amplifying the amplitude of the eye-gaze cue, we effectively triggered gaze-following behavior, which in turn biased the hit probability towards the congruent condition.

**Figure 4.**
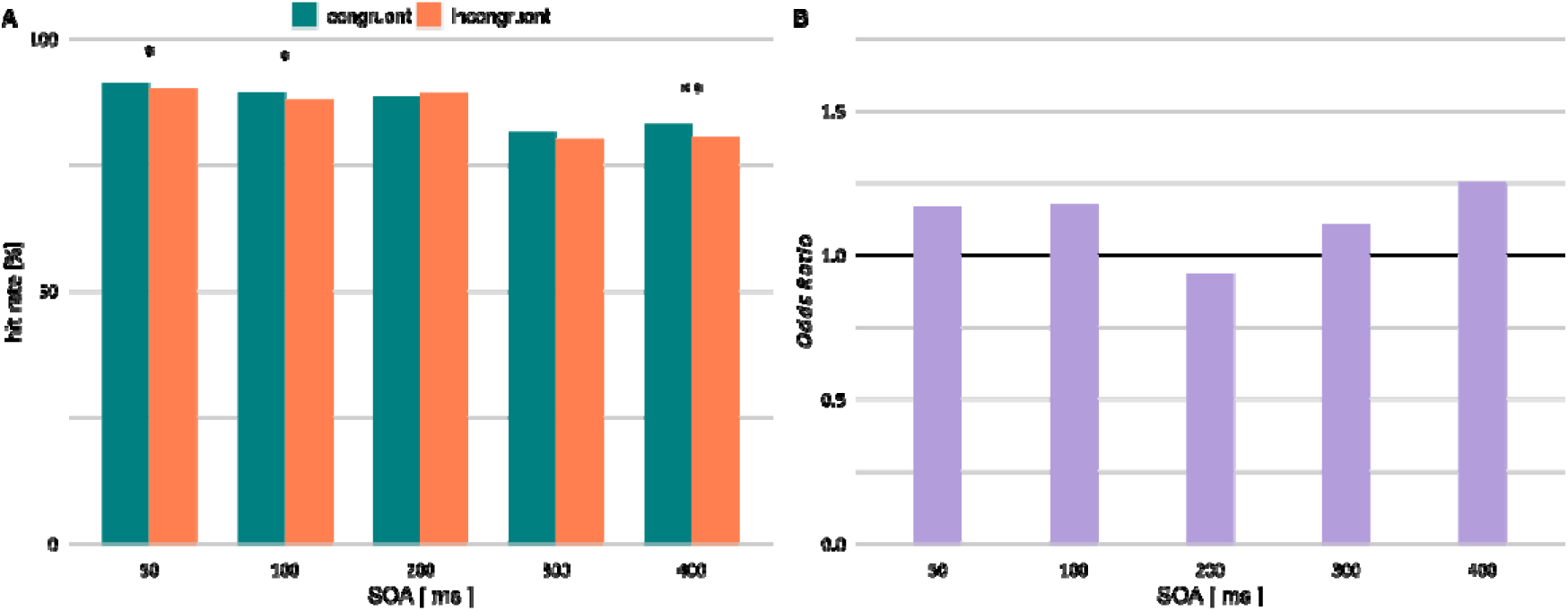
hit rate and OR as a function of SOA for the exaggerated eye-only gaze cue in monkey E. *: p<0.05; **: p<0.01; ***: p<0.001.

**Table 4.** presents the GLMM and ANOVA outputs for the fixed predictors in the eyes-only gaze condition with large amplitude.

|  | monkey E |  |
| --- | --- | --- |
|  | *Wald $\chi^2$* | *log-odds* |
| intercept | 30.46 *** | 0.58 *** |
| congruency (ref: incongruent) | 5.00 *** | -0.16 * |
| SOA (ref: 50) | 72.41 *** | - |
| 100 | - | -0.19 |
| 200 | - | 0.84 *** |
| 300 | - | -0.10 |
| 400 | - | 0.93 *** |
| target location (ref: right) | 250.29 *** | 0.29 *** |
| normalized log RT | 0.01 | 0.01 |
| SOA x congruency | 10.93 | - |
| 100:congruency | - | -0.01 |
| 200:congruency | - | 0.22 * |
| 300:congruency | - | 0.05 |
| 400:congruency | - | -0.07 |
| SOA x target location | 588.86 *** | - |
| 100:target location | - | -0.02 |
| 200:target location | - | -0.37 *** |
| 300:target location | - | -0.22 *** |
| 400:target location | - | -0.47 *** |
| SOA x normalized log RT | 31.49 ** | - |
| 100: normalized log RT | - | 0.16 * |
| 200: normalized log RT | - | 0.25 *** |
| 300: normalized log RT | - | 0.25 *** |
| 400: normalized log RT | - | 0.29 *** |
| congruency x normalized log RT | 0.28 | 0.02 |

number of observations: 25926; random predictor (session number = 28): variance = 0.070; adjusted ICC = 0.020 \* p<0.05 \*\* p<0.01 \*\*\* p<0.001

## Discussion

While previous work has firmly established that rhesus monkeys use conspecifics’ head-gaze to shift their focus of attention, either overtly or covertly, towards the otheŕs object of interest, the role of the otherś eye gaze has remained contentious. Hence, in this experiment, we aimed to address whether object-oriented gaze cues demonstrated by a conspecific’s eyes in the absence of accompanying head turns can guide covert shifts in visual spatial attention in rhesus monkeys. Rather than relying on videos or static images of real conspecifics performing gaze shifts, we used a realistic rhesus monkey head avatar to display object-directed eye gaze, presented either with or without accompanying head turns. The monkey observers were then tasked with detecting near-threshold luminance changes at one of two possible visual targets, with the demonstratoŕs gaze directed toward the target in 50% of trials. Our results revealed significantly higher accuracy when the demonstratoŕs gaze was directed toward the target object, but this effect occurred only when the gaze shift involved head turns. Eye-gaze shifts of identical amplitude in the absence of a head-movement turned out to be ineffective. However, large-amplitude eye-gaze shifts, when presented at a closer distance simulating near-field social interactions, successfully enhanced luminance change detection, indicating covert shifts of attention. These findings suggest that rhesus monkeys are in principle capable of exploiting the spatial information offered by a conspecifićs eyes. Yet, this ability seems to be primarily engaged under close-range conditions, likely suitable to facilitate the scrutiny of the direction of a conspecifićs eyes which are arguably less conspicuous than the eyes of humans. Thus, poorer input quality, rather than a fundamental absence of the necessary perceptual capacity, appear to limit rhesus monkeys’ ability to respond to isolated eye-gaze cues.

To investigate the influence of gaze cues preceding the luminance change on detection accuracy, we compared performance in congruent and incongruent conditions, i.e., trials in which the demonstrator’s gaze was directed toward the target object undergoing a luminance change, versus trials in which the gaze was directed toward the distractor. We hypothesized that higher detection accuracy in congruent trials, relative to incongruent trials, would reflect a covert shift of attention in the observer monkey, guided by the gaze cues presented by the avatar. In incongruent trials, the luminance change was expected to be more frequently missed due to attentional misallocation to the opposite target. A strict central fixation requirement during gaze cue presentation precluded overt attentional shifts via eye movements. Instead of indicating the demonstratoŕs gaze direction, the observers had learned to use saccades to indicate the object they judged to have undergone a luminance change. As our experimental design eliminated any reward-based incentive for gaze-following, any performance enhancements in congruent trials would reflect spontaneous covert gaze-following tendencies, persisting despite the absence of obvious benefits for the observer.

The observed gaze-mediated attentional modulation occurred as early as 50ms relative to the gaze cue, no matter if the gaze shift was based on a head turn or an eye movement from the demonstrator brought into closer view with larger amplitude. Such rapid attentional shifts are consistent with a reflexive mechanism that relies on a few conspicuous exogenous features that may be captured without the need for extensive and therefore time-consuming processing^74–76^. This early effect is supported by substantial evidence from studies demonstrating that neural correlates of covert attentional shifts can be detected in the thalamus and early visual cortical areas as early as 50 ms following cue presentation in non-human primates using Posner-type paradigms^74,77,78^. This effect also aligns with the behavioral findings from a previous report showing that rhesus monkeys covertly shift their attention to targets indicated by conspecificś head-gaze after similarly short latencies^31^. In this study the observer was asked to detect a luminance change of a randomly selected spatial target, to be identified either employing spatial information offered by the otheŕs head gaze or, alternatively the otheŕs identity based on a learned association between identity and target position. In case the observer was instructed to follow the demonstratoŕs gaze, a deployment of attention towards the gaze-congruent target was found to start as early as 50 ms and persisted for up to 500 ms. When the prevailing rule required to ignore the otheŕs gaze and to rely on the otheŕs identity to identify the target, attention was shifted comparatively sustained to the identity-congruent target, yet the shift started significantly later, after only about 300 ms. However, in the preceding period starting around 50 msec an early, transient shift of attention toward the gaze -congruent (but incorrect) target persisted, compatible with limited executive control of an inappropriate shift of attention during this initial period. These results suggested that gaze-driven shifts of covert attention consist of two components: an early, largely automatic component that kicks in too fast to be prone to top-down cognitive control, and a later component, modifiable by control signals available only after time-consuming processing.

The early deployment of covert attentional shifts observed in the present study contrasts with the findings of Deaner and Platt^21^, who reported that reflexive, gaze-driven covert shifts of attention did not occur earlier than 200 ms following the presentation of a gaze cue. Aside from the current work, their study remains the only one to directly investigate the influence of conspecifics’ eyes, either with or without accompanying head turns demonstrating nonpredictive gaze cues on covert attentional shifts in rhesus monkeys. In their paradigm, static, greyscale images of a monkey face with averted eye or head-gaze were shown for varying durations (100, 200, 400, and 800 ms), following the disappearance of a central fixation square. Monkeys were then required to make a saccade to a target appearing at either a gaze-congruent or incongruent location. Although their findings generally support the notion that rhesus monkeys can use conspecific eye or head-gaze direction to covertly shift attention, the late onset of the shift seems not easily reconcilable with the demonstration of an early gaze-dependent attention reflex by Marciniak’s and the present study. A possible explanation may be provided by key methodological differences. Whereas Deaner and Platt presented gaze cues in isolation, without a preceding reference image, both our study and that of Marciniak used a two-step cueing approach: a forward-looking face with direct gaze preceded the gaze cue. We propose that this well-defined transition between direct and averted gaze may render the change in gaze direction, precisely located in time, more salient and easier to detect, thereby accelerating the recognition of behaviorally relevant visual features. Conversely, when a gaze cue is presented without prior direct gaze serving as a baseline, the extraction of critical spatial-cueing information may be delayed. This interpretation is supported by findings from two previous behavioral studies, one in humans and the other in rhesus macaques, which reported a significantly stronger gaze-cueing effect when the cue was preceded by a centrally presented, forward-looking face, compared to trials without such a referential stimulus^43,79^.

A key finding of the present study was the demonstration of covert visual attentional modulation in response to isolated eye-gaze cues, effectively rejecting the hypothesis that rhesus macaques may lack the perceptual capacity to use spatial information provided by the eyes to guide the allocation of attention. Instead, our results highlight the critical influence of two crucial factors in eliciting gaze-following behavior: the distance between the demonstrator and the observer, and the amplitude of the eye-gaze cue. When the eye-gaze cue is subtle and observed from a greater distance, it may go unnoticed. In contrast, when the cue is viewed from close proximity and is sufficiently large in amplitude, it effectively elicits gaze-following behavior in rhesus monkeys. Because we simultaneously manipulated both proximity and amplitude, it is difficult to determine which factor had the greater impact. In any case, our findings clearly indicate that the ability to follow eye gaze is present in rhesus monkeys, a species whose evolutionary lineage diverged from that of modern humans more than 30 million years ago^80^. This suggests that the capability to use eye gaze direction for extracting behaviorally relevant information dates back at least to the last common ancestor of old word monkeys (which include rhesus monkeys) and hominids, i.e. apes and both modern and pre-modern humans. This behavioral finding is in line with neurophysiological data from single-unit recordings in rhesus monkeys, which have identified neurons in the superior temporal sulcus (STS) that are selectively tuned to the direction of eye gaze, independent of head orientation^81,82^. However, it remains unclear to what extent this ability is employed in everyday social interactions, or how effectively eye-gaze cues function for long-range communication in this species.

The prevailing view holds that humans possess a markedly superior capacity for gaze following relative to non-human primates (NHPs). This claim partly rests on a series of studies reporting an absence of eye-gaze following in NHPs. So far, eleven investigations have assessed eye-gaze following in NHPs^19,21,23,27,33–35,42–44,53^, and five of these reported a complete absence of eye-gaze following^19,23,33,44^ or weaker eye-gaze following compared to head-gaze^42^. Notably, all five employed live human demonstrators, but it is not clear whether human gaze cues constitute ecologically valid stimuli for NHPs. Furthermore, failure to respond to human cues does not necessarily imply a general incapacity for following the eye-gaze cues of conspecifics in NHPs. In contrast to these null findings, a few studies using the same live-human demonstrator paradigm have reported robust eye gaze-following responses in NHPs^27,34,35^. Such inconsistency may stem from uncontrolled variables, such as the quantity and quality of exposure to human caregivers or experimenters and from age-related differences in gaze-following propensity. Indeed, increased exposure to humans correlates positively with NHPs’ ability to follow human eye-gaze^20,23,24,32,33^, whereas the influence of age on this capacity remains unclear. Some reports document improvements in gaze following with age^23,24^; for example, Ferrari et al. (2000)^23^ demonstrated that adult pig-tailed macaques (*Macaca nemestrina*) follow both head- and eye-gaze cues, whereas juveniles rely solely on head-gaze. Contrary to these reports, Tomasello et al. (2007)^42^ and Parron & Meguerditchian (2016)^32^ observed the opposite pattern. Additional factors, such as duration of exposure to human gaze cues and rates of habituation may covary with age, further complicating interpretations of age-related effects^32^. Moreover, all studies yielding negative results lacked high-resolution eye-tracking and instead usually relied on video-recorded sessions scored by human observers^19,23,33,42,44^. Standard video cameras operating at 25–30 frames per second used in these studies often miss or distort (i.e., “motion blur” effect) very rapid movement like saccades driven by overt shifts of visual attention in response to gaze cues. Additionally, some paradigms required NHPs to follow a human experimenter’s gaze into empty space^23,42^. Repeated exposure to situations where gaze following yields no clear target can induce habituation, attenuating subsequent gaze-following responses^32,42^ as these behaviors are typically object-directed rather than spatially indiscriminate. Only one ‘‘negative’’ study directly contrasted congruent and incongruent gaze cues, yet surprisingly each subject experienced only a single trial per condition. Collectively, the evidence supporting the notion that eye-gaze following is unique to humans is undermined by substantial methodological limitations, casting doubt on the reliability of the conclusion that NHPs lack this capacity.

Given that the lack of sound empirical evidence supporting the claim that eye gaze-following behavior is uniquely human, it is crucial to reconsider the other main argument supporting exclusivity of eye-gaze following in humans: the supposed exceptional visibility of human eyes. Early qualitative studies by Kobayashi and Kohshima (1997, 2001)^47,48^, building on Morris (1985)^83^, argued that humans possess uniquely visible eyes, characterized by a pale sclera, a high eye width-to-height ratio (WHR), and substantial scleral exposure. However, these conclusions have been challenged by recent findings on several grounds. First, pale peri-iridal tissues have been documented in various Old World and New World monkey species, and significant variation exists within species concerning scleral pigmentation and eye shape^49,84–89^. For instance, 17% of Ngogo chimpanzees, despite typically having dark sclera, exhibit pale sclera^87^, while many humans living in equatorial or rural regions commonly have conjunctival pigmentation^90–95^. Additionally, WHR varies across humans, and NHPs like orangutans have WHR values comparable to humans^29^. Second, gaze visibility is determined by the contrast between iris, sclera, and peri-orbital skin rather than scleral brightness alone, with studies revealing no significant luminance contrast differences among humans, chimpanzees, and bonobos^85,96^. Third, early investigations focused on direct gaze when comparing the area of exposed sclera across species, while subsequent studies have shown that, when gaze is averted, the proportion of visible sclera does not differ significantly between humans and gorillas^49^. The contradictory evidence undermines the hypothesis that humans possess uniquely conspicuous ocular morphology. Consequently, it challenges the claim that this trait underpins superior gaze-following capabilities, and exposes the limitations of the initial, overly simplistic visibility metrics. Hence, a more nuanced view should be adopted because gaze-cue visibility likely depends on many factors some of which may be interacting such as observer-demonstrator distance, ambient illumination, and the combined hue-luminance contrasts of the iris and sclera. Moreover, the specific contribution of hue contrast among ocular structures remains unexplored; configurations matched for luminance may still differ in hue contrast and, therefore, in perceptual salience.

In summary current evidence, including our own findings, rejects a qualitative gap in eye gaze-following ability between humans and non-human primates (NHPs). However, nevertheless there could still be significant quantitative differences. Three comparative studies have directly addressed this question. Two reported markedly larger effects in humans: Tomasello et al. (2007)^42^ found that 12- to 18-month-old infants showed 1.26-fold higher gaze-following accuracy than great apes (Cohen’s d = 1.53 vs. 1.22, respectively), and Deaner & Platt (2003)^21^ observed reaction-time and eye-position effects in humans that were 1.7–3.6 times those of monkeys. Conversely, Itakura & Tanaka (1998)^27^ detected no species difference in an object-choice task involving chimpanzees, an orangutan, and 23-month-old infants. This limited, methodologically diverse literature tentatively favors greater human proficiency but precludes firm conclusions, highlighting the need for more well-controlled experiments. Such comparative studies should present gaze cues from both human and non-human conspecifics enabling a more precise dissection of intra- and inter-specific similarities in gaze-following performance. High-fidelity virtual avatars, like those we incorporated in the present study, provide an optimal platform for this line of research. They enable systematic manipulation of factors influencing gaze cue salience, including demonstrator–observer distance, illumination, scleral and peri-orbital luminance and hue contrasts, eye shape, exposed scleral area, and gaze-motion kinematics, while maintaining ecologically valid stimuli.

In conclusion, the present study demonstrates that the visuomotor system of rhesus monkeys is capable of following eye gaze, provided the gaze cue is sufficiently salient, such as when viewed at close proximity with large enough amplitude. The gaze-driven modulation of covert visual spatial attention occurs reflexively in response to salient visual input. Notably, we also observed attentional effects persisting at longer cue-target intervals, indicating the possible engagement of endogenous, goal-directed attention processes in gaze-following. Furthermore, the use of a realistic monkey avatar proved essential for addressing our research question with greater precision which opens promising avenues for future studies exploring the sensory and cognitive mechanisms underlying social cue processing in nonhuman primates and humans.

## Data and software availability statement

The data has been deposited on the Open Science Framework (OSF) and is available upon request.

Those scripts used for the statistical analysis of the data are publicly available at: https://github.com/shafiei-masih/SocialAttention_monkeys

## References

1. Mehrabian, A. (2017). Nonverbal Communication (Routledge) 10.4324/9781351308724.

2. Drijvers, L., and Holler, J. (2023). The multimodal facilitation effect in human communication. Psychon Bull Rev 30, 792–801. 10.3758/s13423-022-02178-x.

3. Morioka, S., Osumi, M., Shiotani, M., Nobusako, S., Maeoka, H., Okada, Y., Hiyamizu, M., and Matsuo, A. (2016). Incongruence between Verbal and Non-Verbal Information Enhances the Late Positive Potential. PLoS One 11, e0164633. 10.1371/journal.pone.0164633.

4. Bock, S.W., Dicke, P., and Thier, P. (2008). How precise is gaze following in humans? Vision Research 48, 946–957. 10.1016/j.visres.2008.01.011.

5. Butterworth, G., and Jarrett, N. (1991). What minds have in common is space: Spatial mechanisms serving joint visual attention in infancy. British Journal of Developmental Psychology 9, 55–72. 10.1111/j.2044-835X.1991.tb00862.x.

6. Charman, T., Baron-Cohen, S., Swettenham, J., Baird, G., Cox, A., and Drew, A. (2000). Testing joint attention, imitation, and play as infancy precursors to language and theory of mind. Cognitive Development 15, 481–498. 10.1016/S0885-2014(01)00037-5.

7. Sodian, B., and Kristen-Antonow, S. (2015). Declarative joint attention as a foundation of theory of mind. Dev Psychol 51, 1190–1200. 10.1037/dev0000039.

8. Morales, M., Mundy, P., and Rojas, J. (1998). Following the direction of gaze and language development in 6-month-olds. Infant Behavior and Development 21, 373–377. 10.1016/S0163-6383(98)90014-5.

9. Brooks, R., and Meltzoff, A.N. (2008). Infant gaze following and pointing predict accelerated vocabulary growth through two years of age: a longitudinal, growth curve modeling study. Journal of Child Language 35, 207–220. 10.1017/S030500090700829X.

10. Beuker, K.T., Rommelse, N.N.J., Donders, R., and Buitelaar, J.K. (2013). Development of early communication skills in the first two years of life. Infant Behav Dev 36, 71–83. 10.1016/j.infbeh.2012.11.001.

11. Dawson, G., Toth, K., Abbott, R., Osterling, J., Munson, J., Estes, A., and Liaw, J. (2004). Early social attention impairments in autism: social orienting, joint attention, and attention to distress. Dev Psychol 40, 271–283. 10.1037/0012-1649.40.2.271.

12. Elsabbagh, M., Mercure, E., Hudry, K., Chandler, S., Pasco, G., Charman, T., Pickles, A., Baron-Cohen, S., Bolton, P., Johnson, M.H., et al. (2012). Infant neural sensitivity to dynamic eye gaze is associated with later emerging autism. Curr Biol 22, 338–342. 10.1016/j.cub.2011.12.056.

13. Thorup, E., Nyström, P., Gredebäck, G., Bölte, S., Falck-Ytter, T., and EASE Team (2016). Altered gaze following during live interaction in infants at risk for autism: an eye tracking study. Mol Autism 7, 12. 10.1186/s13229-016-0069-9.

14. Congiu, S., Fadda, R., Doneddu, G., and Striano, T. (2016). Impaired representational gaze following in children with autism spectrum disorder. Res Dev Disabil 57, 11–17. 10.1016/j.ridd.2016.06.008.

15. Cheney, D., Seyfarth, R., and Smuts, B. (1986). Social Relationships and Social Cognition in Nonhuman Primates. Science 234, 1361–1366. 10.1126/science.3538419.

16. Platt, M.L., Seyfarth, R.M., and Cheney, D.L. (2016). Adaptations for social cognition in the primate brain. Philos Trans R Soc Lond B Biol Sci 371, 20150096. 10.1098/rstb.2015.0096.

17. Anderson, J.R., and Mitchell, R.W. (1999). Macaques but Not Lemurs Co-Orient Visually with Humans. IJFP 70, 17–22. 10.1159/000021670.

18. Bräuer, J., Call, J., and Tomasello, M. (2005). All Great Ape Species Follow Gaze to Distant Locations and Around Barriers. Journal of Comparative Psychology 119, 145–154. 10.1037/0735-7036.119.2.145.

19. Burkart, J., and Heschl, A. (2006). Geometrical gaze following in common marmosets (Callithrix jacchus). Journal of Comparative Psychology 120, 120–130. 10.1037/0735-7036.120.2.120.

20. Byrnit, J.T. (2009). Gorillas’ (Gorilla gorilla) use of experimenter-given manual and facial cues in an object-choice task. Anim Cogn 12, 401–404. 10.1007/s10071-008-0200-1.

21. Deaner, R.O., and Platt, M.L. (2003). Reflexive Social Attention in Monkeys and Humans. Current Biology 13, 1609–1613. 10.1016/j.cub.2003.08.025.

22. Emery, N.J., Lorincz, E.N., Perrett, D.I., Oram, M.W., and Baker, C.I. (1997). Gaze Following and Joint Attention in Rhesus Monkeys (Macaca mulatto). Journal of Comparative Psychology 111, 286–293.

23. Ferrari, P.F., Kohler, E., Fogassi, L., and Gallese, V. (2000). The ability to follow eye gaze and its emergence during development in macaque monkeys. Proceedings of the National Academy of Sciences 97, 13997–14002. 10.1073/pnas.250241197.

24. Ferrari, P.F., Coude, G., Gallese, V., and Fogassi, L. (2008). Having access to others’ mind through gaze: the role of ontogenetic and learning processes in gaze-following behavior of macaques. Soc Neurosci 3, 239–249. 10.1080/17470910701429065.

25. Goossens, B.M.A., Dekleva, M., Reader, S.M., Sterck, E.H.M., and Bolhuis, J.J. (2008). Gaze following in monkeys is modulated by observed facial expressions. Animal Behaviour 75, 1673–1681. 10.1016/j.anbehav.2007.10.020.

26. Itakura, S. (1996). An exploratory study of gaze-monitoring in nonhuman primates. Japanese Psychological Research 38, 174–180. 10.1111/j.1468-5884.1996.tb00022.x.

27. Itakura, S., and Tanaka, M. (1998). Use of experimenter-given cues during object-choice tasks by chimpanzees (Pan troglodytes), an orangutan (Pongo pygmaeus), and human infants (Homo sapiens). Journal of Comparative Psychology 112, 119–126. 10.1037/0735-7036.112.2.119.

28. Kano, F., and Call, J. (2014). Cross-species variation in gaze following and conspecific preference among great apes, human infants and adults. Animal Behaviour 91, 137–150. 10.1016/j.anbehav.2014.03.011.

29. Kaplan, G., and Rogers, L.J. (2002). Patterns of Gazing in Orangutans (Pongo pygmaeus). International Journal of Primatology 23, 501–526. 10.1023/A:1014913532057.

30. Liebal, K., and Kaminski, J. (2012). Gibbons (Hylobates pileatus, H. moloch, H. lar, Symphalangus syndactylus) follow human gaze, but do not take the visual perspective of others. Anim Cogn 15, 1211–1216. 10.1007/s10071-012-0543-5.

31. Marciniak, K., Dicke, P.W., and Thier, P. (2015). Monkeys head-gaze following is fast, precise and not fully suppressible. Proceedings of the Royal Society B: Biological Sciences 282, 20151020. 10.1098/rspb.2015.1020.

32. Parron, C., and Meguerditchian, A. (2016). Gaze following in baboons (Papio anubis): juveniles adjust their gaze and body position to human’s head redirections. Am J Primatol 78, 1265–1271. 10.1002/ajp.22580.

33. Peignot, P., and Anderson, J.R. (1999). Use of experimenter-given manual and facial cues by gorillas (Gorilla gorilla) in an object-choice task. Journal of Comparative Psychology 113, 253–260. 10.1037/0735-7036.113.3.253.

34. Povinelli, D.J., and Eddy, T.J. (1996). Chimpanzees: Joint Visual Attention. Psychol Sci 7, 129–135. 10.1111/j.1467-9280.1996.tb00345.x.

35. Povinelli, D.J., and Eddy, T.J. (1997). Specificity of gaze-following in young chimpanzees. British Journal of Developmental Psychology 15, 213–222. 10.1111/j.2044-835X.1997.tb00735.x.

36. Rosati, A.G., Arre, A.M., Platt, M.L., and Santos, L.R. (2016). Rhesus monkeys show human-like changes in gaze following across the lifespan. Proceedings of the Royal Society B: Biological Sciences 283, 20160376. 10.1098/rspb.2016.0376.

37. Ruiz, A., Gómez, J.C., Roeder, J.J., and Byrne, R.W. (2009). Gaze following and gaze priming in lemurs. Anim Cogn 12, 427–434. 10.1007/s10071-008-0202-z.

38. Shepherd, S.V., and Platt, M.L. (2008). Spontaneous social orienting and gaze following in ringtailed lemurs (Lemur catta). Anim Cogn 11, 13–20. 10.1007/s10071-007-0083-6.

39. Teufel, C., Gutmann, A., Pirow, R., and Fischer, J. (2010). Facial expressions modulate the ontogenetic trajectory of gaze-following among monkeys. Developmental Science 13, 913–922. 10.1111/j.1467-7687.2010.00956.x.

40. Tomasello, M., Call, J., and Hare, B. (1998). Five primate species follow the visual gaze of conspecifics. Animal Behaviour 55, 1063–1069. 10.1006/anbe.1997.0636.

41. Tomasello, M., Hare, B., and Agnetta, B. (1999). Chimpanzees, *Pan troglodytes*, follow gaze direction geometrically. Animal Behaviour 58, 769–777. 10.1006/anbe.1999.1192.

42. Tomasello, M., Hare, B., Lehmann, H., and Call, J. (2007). Reliance on head versus eyes in the gaze following of great apes and human infants: the cooperative eye hypothesis. Journal of Human Evolution 52, 314–320. 10.1016/j.jhevol.2006.10.001.

43. Yu, D., Teichert, T., and Ferrera, V.P. (2012). Orienting of Attention to Gaze Direction Cues in Rhesus Macaques: Species-Specificity, and Effects of Cue Motion and Reward Predictiveness. Front Psychol 3, 202. 10.3389/fpsyg.2012.00202.

44. Santos, L.R., and Hauser, M.D. (1999). How monkeys see the eyes: cotton-top tamarins’ reaction to changes in visual attention and action. Anim Cogn 2, 131–139. 10.1007/s100710050033.

45. Moors, P., Germeys, F., Pomianowska, I., and Verfaillie, K. (2015). Perceiving where another person is looking: the integration of head and body information in estimating another person’s gaze. Front Psychol 6, 909. 10.3389/fpsyg.2015.00909.

46. Atabaki, A., Marciniak, K., Dicke, P.W., and Thier, P. (2015). Assessing the precision of gaze following using a stereoscopic 3D virtual reality setting. Vision Res 112, 68–82. 10.1016/j.visres.2015.04.015.

47. Kobayashi, H., and Kohshima, S. (2001). Unique morphology of the human eye and its adaptive meaning: comparative studies on external morphology of the primate eye. Journal of Human Evolution 40, 419–435. 10.1006/JHEV.2001.0468.

48. Kobayashi, H., and Kohshima, S. (1997). Unique morphology of the human eye. Nature 1997 387:6635 387, 767–768. 10.1038/42842.

49. Mayhew, J.A., and Gómez, J.-C. (2015). Gorillas with white sclera: A naturally occurring variation in a morphological trait linked to social cognitive functions. American Journal of Primatology 77, 869–877. 10.1002/ajp.22411.

50. Perea-García, J.O., Kret, M.E., Monteiro, A., and Hobaiter, C. (2019). Scleral pigmentation leads to conspicuous, not cryptic, eye morphology in chimpanzees. Proceedings of the National Academy of Sciences of the United States of America 116, 19248–19250. 10.1073/PNAS.1911410116/ASSET/1A31A704-262B-409C-9524-0F607738201B/ASSETS/GRAPHIC/PNAS.1911410116FIG02.JPEG.

51. Mearing, A.S., and Koops, K. (2021). Quantifying gaze conspicuousness: Are humans distinct from chimpanzees and bonobos? Journal of Human Evolution 157, 103043. 10.1016/J.JHEVOL.2021.103043.

52. Perea-García, J.O., Teuben, A., and Caspar, K.R. (2025). Look past the cooperative eye hypothesis: reconsidering the evolution of human eye appearance. Biological Reviews n/a. 10.1111/brv.70033.

53. Lorincz, E.N., Baker, C.I., and Perrett, D.I. (2001). Visual cues for attention following in rhesus monkeys. In Picture Perception in Animals (Routledge).

54. Tomasello, M., Hare, B., and Fogleman, T. (2001). The ontogeny of gaze following in chimpanzees, *Pan troglodytes*, and rhesus macaques, *Macaca mulatta*. Animal Behaviour 61, 335–343. 10.1006/anbe.2000.1598.

55. Siebert, R., Taubert, N., Spadacenta, S., Dicke, P.W., Giese, M.A., and Thier, P. (2020). A Naturalistic Dynamic Monkey Head Avatar Elicits Species-Typical Reactions and Overcomes the Uncanny Valley. eNeuro 7. 10.1523/ENEURO.0524-19.2020.

56. Lu, X., Wang, Q., Li, X., Wang, G., Chen, Y., Li, X., and Li, H. (2023). Connectivity reveals homology between the visual systems of the human and macaque brains. Front. Neurosci. 17. 10.3389/fnins.2023.1207340.

57. Prsa, M., and Thier, P. (2011). The role of the cerebellum in saccadic adaptation as a window into neural mechanisms of motor learning. Eur J Neurosci 33, 2114–2128. 10.1111/j.1460-9568.2011.07693.x.

58. Bechert, K., and Koenig, E. (1996). A search coil system with automatic field stabilization, calibration, and geometric processing for eye movement recording in humans. Neuro-Ophthalmology. 10.3109/01658109609009677.

59. R Core Team (2024). R: A Language and Environment for Statistical Computing (R Foundation for Statistical Computing).

60. Wickham, H. (2016). ggplot2: Elegant Graphics for Data Analysis (Springer-Verlag New York).

61. Bates, D., Mächler, M., Bolker, B., and Walker, S. (2015). Fitting Linear Mixed-Effects Models Using lme4. Journal of Statistical Software 67, 1–48. 10.18637/jss.v067.i01.

62. Fox, John and Weisberg, Sanford (2019). An R Companion to Applied Regression Third. (Sage).

63. Russell V. Lenth (2025). emmeans: Estimated Marginal Means, aka Least-Squares Means. Version 1.10.7.

64. Daniel Lüdecke, Mattan S. Ben-Shachar, Indrajeet Patil, Philip Waggoner, and Dominique Makowski (2021). performance: An R Package for Assessment, Comparison and Testing of Statistical Models. 6, 3139. 10.21105/joss.03139.

65. The MathWorks, Inc. (2023). MATLAB (R2023a). Version R2023a.

66. Fox, J., and Monette, G. (1992). Generalized Collinearity Diagnostics. Journal of the American Statistical Association 87, 178–183. 10.1080/01621459.1992.10475190.

67. Symonds, M.R.E., and Moussalli, A. (2011). A brief guide to model selection, multimodel inference and model averaging in behavioural ecology using Akaike’s information criterion. Behav Ecol Sociobiol 65, 13–21. 10.1007/s00265-010-1037-6.

68. Burnham, K.P., and Anderson, D.R. eds. (2004). Model Selection and Multimodel Inference (Springer) 10.1007/b97636.

69. Nakagawa, S., and Schielzeth, H. (2013). A general and simple method for obtaining R2 from generalized linear mixed-effects models. Methods in Ecology and Evolution 4, 133–142. 10.1111/j.2041-210x.2012.00261.x.

70. Bates, D., Mächler, M., Bolker, B., and Walker, S. (2015). Fitting Linear Mixed-Effects Models Using lme4. Journal of Statistical Software 67, 1–48. 10.18637/jss.v067.i01.

71. Benjamini, Y., and Hochberg, Y. (1995). Controlling the False Discovery Rate: A Practical and Powerful Approach to Multiple Testing. Journal of the Royal Statistical Society: Series B (Methodological) 57, 289–300. 10.1111/j.2517-6161.1995.tb02031.x.

72. Luke, S.G. (2017). Evaluating significance in linear mixed-effects models in R. Behav Res 49, 1494–1502. 10.3758/s13428-016-0809-y.

73. Fox, J. (2015). Applied Regression Analysis and Generalized Linear Models (SAGE Publications).

74. Corbetta, M., and Shulman, G.L. (2002). Control of goal-directed and stimulus-driven attention in the brain. Nat Rev Neurosci 3, 201–215. 10.1038/nrn755.

75. Duncan, J., and Humphreys, G.W. (1989). Visual search and stimulus similarity. Psychological Review 96, 433–458. 10.1037/0033-295X.96.3.433.

76. Posner, Michael I. and Cohen, Yoav (1984). Components of visual orienting. In Attention and performance X: Control of language processes (Erlbaum), pp. 531– 556.

77. McAlonan, K., Cavanaugh, J., and Wurtz, R.H. (2008). Guarding the gateway to cortex with attention in visual thalamus. Nature 456, 391–394. 10.1038/nature07382.

78. McAdams, C.J., and Reid, R.C. (2005). Attention modulates the responses of simple cells in monkey primary visual cortex. J Neurosci 25, 11023–11033. 10.1523/JNEUROSCI.2904-05.2005.

79. Driver IV, J., Davis, G., Ricciardelli, P., Kidd, P., Maxwell, E., and Baron-Cohen, S. (1999). Gaze Perception Triggers Reflexive Visuospatial Orienting. Visual Cognition 6, 509–540. 10.1080/135062899394920.

80. Pozzi, L., Hodgson, J.A., Burrell, A.S., Sterner, K.N., Raaum, R.L., and Disotell, T.R. (2014). Primate phylogenetic relationships and divergence dates inferred from complete mitochondrial genomes. Molecular Phylogenetics and Evolution 75, 165–183. 10.1016/j.ympev.2014.02.023.

81. Perrett, D.I., Smith, P. a. J., Potter, D.D., Mistlin, A.J., Head, A.S., Milner, A.D., Jeeves, M.A., and Weiskrantz, L. (1985). Visual cells in the temporal cortex sensitive to face view and gaze direction. Proceedings of the Royal Society of London. Series B. Biological Sciences 223, 293–317. 10.1098/rspb.1985.0003.

82. Yang, Z., and Freiwald, W.A. (2021). Joint encoding of facial identity, orientation, gaze, and expression in the middle dorsal face area. Proceedings of the National Academy of Sciences 118, e2108283118. 10.1073/pnas.2108283118.

83. Morris, Desmond. (1985). Bodywatching: A Field Guide to the Human Species (Crown).

84. Perea García, J.O. (2016). Quantifying ocular morphologies in extant primates for reliable interspecific comparisons. Journal of Language Evolution 1, 151–158. 10.1093/jole/lzw004.

85. Perea-García, J.O., Kret, M.E., Monteiro, A., and Hobaiter, C. (2019). Scleral pigmentation leads to conspicuous, not cryptic, eye morphology in chimpanzees. Proceedings of the National Academy of Sciences 116, 19248–19250. 10.1073/pnas.1911410116.

86. Caspar, K.R., Biggemann, M., Geissmann, T., and Begall, S. (2021). Ocular pigmentation in humans, great apes, and gibbons is not suggestive of communicative functions. Sci Rep 11, 12994. 10.1038/s41598-021-92348-z.

87. Clark, I.R., Lee, K.C., Poux, T., Langergraber, K.E., Mitani, J.C., Watts, D., Reed, J., and Sandel, A.A. (2023). White sclera is present in chimpanzees and other mammals. Journal of Human Evolution 176, 103322. 10.1016/j.jhevol.2022.103322.

88. Perea-García, J.O., Ramarajan, K., Kret, M.E., Hobaiter, C., and Monteiro, A. (2022). Ecological factors are likely drivers of eye shape and colour pattern variations across anthropoid primates. Sci Rep 12, 17240. 10.1038/s41598-022-20900-6.

89. Perea-García, J.O., Massen, J.J.M., Ostner, J., Schülke, O., Castellano-Navarro, A., Gazagne, E., José-Domínguez, J.M., Beltrán-Francés, V., Kaburu, S., Ruppert, N., et al. (2024). Photoregulatory functions drive variation in eye coloration across macaque species. Sci Rep 14, 29115. 10.1038/s41598-024-80643-4.

90. Whittington, C.P., Bresler, S.C., Simon, C., Shields, C.L., and Patel, R.M. (2024). Melanocytic lesions of the conjunctiva: an up-to-date review. Diagnostic Histopathology 30, 37–59. 10.1016/j.mpdhp.2023.10.005.

91. Mann, I. (1966). Culture, Race, Climate and Eye Disease: An Introduction to the Study of Geographic Ophthalmology (Charles C Thomas).

92. Jakobiec, F.A. (1984). The ultrastructure of conjunctival melanocytic tumors. Trans Am Ophthalmol Soc 82, 599–752.

93. Singh, A.D., Campos, O.E., Rhatigan, R.M., Schulman, J.A., and Misra, R.P. (1998). Conjunctival Melanoma in the Black Population. Survey of Ophthalmology 43, 127–133. 10.1016/S0039-6257(98)00020-4.

94. Jakobiec, F.A. (2016). Conjunctival Primary Acquired Melanosis: Is It Time for a New Terminology? American Journal of Ophthalmology 162, 3–19.e1. 10.1016/j.ajo.2015.11.003.

95. Blake, C.R., Lai, W.W., and Edward, D.P. (2003). Racial and Ethnic Differences in Ocular Anatomy. International Ophthalmology Clinics 43, 9.

96. Mearing, A.S., and Koops, K. (2021). Quantifying gaze conspicuousness: Are humans distinct from chimpanzees and bonobos? Journal of Human Evolution 157, 103043. 10.1016/j.jhevol.2021.103043.

