## Supplemental figure 1 for "Rhesus monkeys use both eye and head gaze to reallocate covert spatial attention facilitating visual perception"

### Supplementary figure

**A**

**B**

luminance change (Go signal)

reward

trial begins

with fixating on fixation dot

response

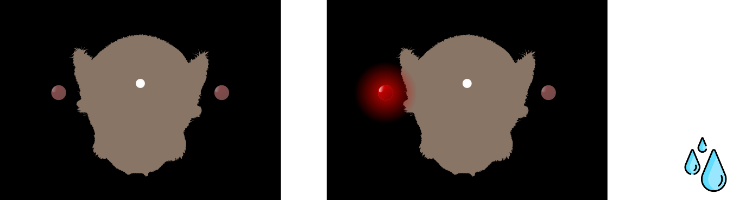

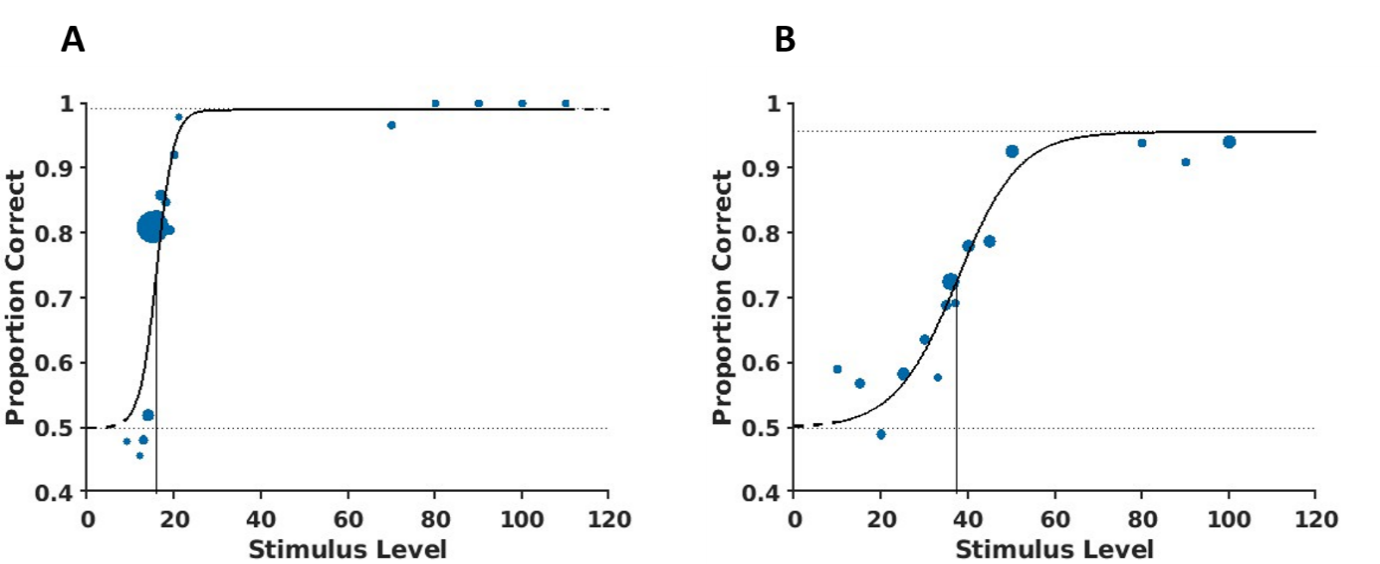

**monkey E**

**monkey C**

**Supplementary figure 1.** A. Schematic of the task used for perceptual threshold assessment. B. Psychometric functions describing the association between the hit rate and luminance change detection for monkey E and C. Stimulus level refers to a total of 512 steps in which the luminance of the LEDs could be varied. The number of trials was 1820 for monkey E and 2218 for monkey C.
